# Disease mediated changes to life history and demography threaten the survival of European amphibian populations

**DOI:** 10.1101/178723

**Authors:** Lewis J. Campbell, Trenton W. J. Garner, Giulia Tessa, Benjamin C. Scheele, Amber G.F. Griffiths, Lena Wilfert, Xavier A. Harrison

## Abstract

Infectious diseases can influence the life history strategy of their hosts and such influences subsequently impact the demography of infected populations, reducing viability independently of increased mortality or morbidity. Amphibians are the most threatened group of vertebrates and emerging infectious diseases play a large role in their population declines. Viruses of genus *Ranavirus* are responsible for one of the deadliest of these diseases. To date no work has evaluated the impact of ranaviruses on host life-history post metamorphosis or population demographic structure at the individual level. In this study, we used skeletochronology and morphology to evaluate the impact of ranaviruses on the demography of populations of European common frog (*Rana temporaria*) in the United Kingdom. We compared ecologically similar populations that differed only in their historical presence or absence of ranaviral disease. Our results suggest that ranaviruses are associated with shifts in the age structure of infected populations, potentially caused by increased adult mortality and associated shifts in the life history of younger age classes. Population projection models indicate that such age truncation could heighten the vulnerability of frog populations to stochastic environmental challenges. Our individual level data provide further compelling evidence that the emergence of infectious diseases can alter host demography, subsequently increasing population vulnerability to additional stressors.

## Introduction

Infectious diseases can have major long-term impacts on host life history strategies, often leading to subsequent shifts in the demographic structure of host populations [1–4]. Within age-structured populations, this is primarily caused by compensatory changes in the vital rates (rates of growth, fecundity and survival) of younger age classes that occur in response to high levels of extrinsic adult mortality (death of adult animals attributable to external factors such as disease or predation [5,6]). In order to maximise individual fitness within an environment of high extrinsic adult mortality, selection for increased juvenile survival and developmental rates, decreased size and age at sexual maturity and ultimately an increased adult life span (deceased intrinsic adult mortality) will occur [7]. This has been empirically borne out in a number of systems in response to several sources of mortality including; predation [8], over-harvesting [9] and experimentally induced adult mortality [7].

Impacts on population demographic structure potentially brought about by changes to individual life history strategies can have substantial effects on the growth and stability of infected populations [10]. This has recently been well demonstrated by Scheele *et al.* [11] who observed that infection with the lethal fungal pathogen *Batrachochytrium dendrobatidis* (*Bd*) causes high levels of adult mortality in the endangered Australian alpine tree frog (*Litoria verreauxii alpina*). High adult mortality resulted in a severe truncation of the age structure of infected populations. Subsequent population projection modelling showed that such *Bd* positive populations are much more vulnerable to decline due to stochastic recruitment failure caused by a variable environment than *Bd* negative populations [11], highlighting an important, yet relatively unexplored mechanism by which infectious diseases can impact their hosts. Infectious diseases are emerging at a faster rate and threatening a larger range of species than at any prior point in history [12], therefore it is imperative that we aim to better understand such mechanisms, particularly in species of conservation concern.

Amphibians are the most imperilled class of vertebrates [13]. Aside from threats such as habitat loss [14], over-harvesting [15] and climate change [16], one major driver of continued amphibian declines is the emergence of a suite of infectious diseases [17]. One of the most widespread and deadly of these diseases, ranavirosis, is caused by viral pathogens belonging to the genus *Ranavirus* [18,19]. Ranaviruses are globally distributed and infect and kill species from 3 classes of ectothermic vertebrates [20–22]. Clinical ranavirosis is often characterised by severe dermal ulcerations as well as haemorrhaging and lesions affecting the internal organs [22,23]. Outbreaks are known to cause mass mortality events, with up to 90% mortality reported in some instances [19]. Evidence from the UK shows that emergence within a population can potentially lead to greater than 80% declines in population sizes, followed by suppression or in some cases local extinction [24]. Susceptibility varies among species but a number of ecological risk factors that contribute to species susceptibility, including life history strategy, have been identified [25]. The presence of ranaviruses within a habitat has also been shown to accelerate the developmental rates of larval North American anurans [26]. To date no work has evaluated the impact of ranaviruses on life history strategies post metamorphosis, or on the demographic make-up of populations using individual level data.

In this study, we utilised the unique comparative field system borne out of the Frog Mortality Project (FMP; see [24,27] for details) to study the impacts of ranaviral disease history on demographic structure and individual life history of European common frog (*R. temporaria*) in the United Kingdom. We used a combination of skeletochronology (age determination by skeletal growth rings) and morphometric data collected from wild caught *R. temporaria* to test two hypotheses: i) that the age structure of populations with a positive disease history of ranavirus will be truncated and ii) that frogs originating from ranavirus positive field sites will display reduced body size and more rapid growth compared to their counterparts from ostensibly disease-free populations. Additionally, we performed population matrix modelling to investigate whether persistent ranavirosis heightens population susceptibility to additional external stressors.

## Methods

### Field Sampling

Potential field sites were drawn from the FMP database of *R. temporaria* populations known to have experienced at least one mass mortality event due to ranavirosis and a complimentary database of putatively disease free *R. temporaria* populations (see [24] for more detailed selection criteria). Five populations of each disease history were selected (Fig 1). Field sites were attended during the spring breeding season and each was sampled for one day. Sampling involved searching for frogs within and around the proximity of the breeding ponds. As many individuals as possible, including any amplexing pairs, were located and placed into plastic holding tanks (care was taken not to cause paired frogs to release). Since as many frogs as possible were captured at each field site by the same individual researcher, sampling effort is considered to be approximately equal. As all frogs were found to be above the age minimum age of sexual maturity, and juvenile anurans rarely return to breeding ponds [28,29, but see 30], all were assumed to be members of the breeding population.

**Fig 1.**
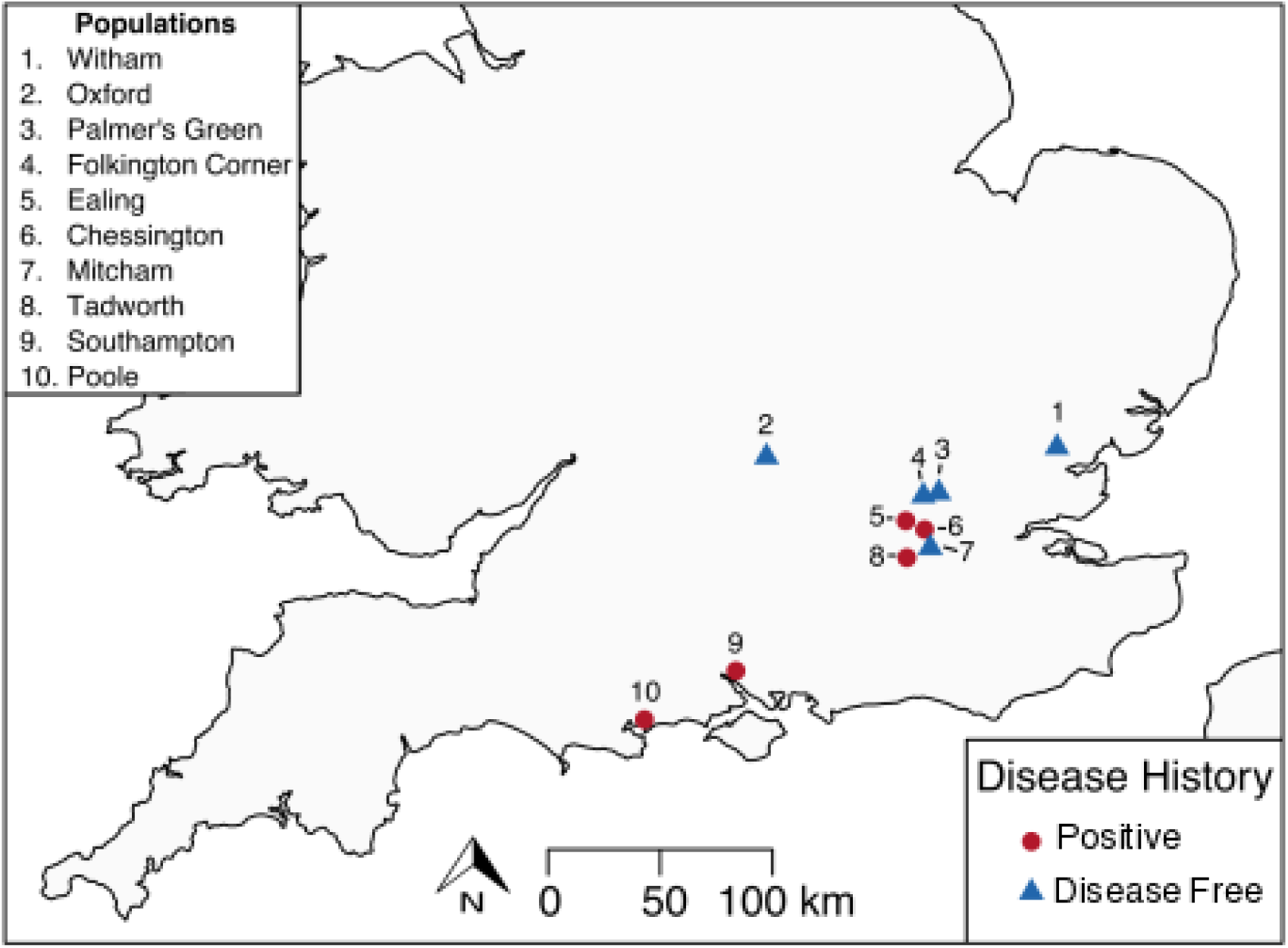
Map of the locations of sampled populations within the southern United Kingdom. Field sites were drawn from the Frog Mortality Project database of populations known to have experienced FV3 mass mortality events and a complimentary database of populations known to have been FV3 free since disease emergence in the early 1990s.

Snout to vent length was measured using 0.1mm scale callipers. For unpaired frogs, body weight was recorded using a 100g maximum weight (1g increments) Pesola drop scale. Frogs were weighed in a plastic zip lock bag of known weight (4g), which was subtracted from the total reading of the scale. A clean zip lock bag was used for each individual.

The distal portion of the 1^st^ (inside) digit of a hind limb of each frog was clipped using surgical scissors. To minimise the potential for pain and the possibility of infection, a topical disinfectant that contained an analgesic (Bactine; WellSpring Pharmaceutical, Sarasota, USA) was applied to the surgical area prior to the procedure. Toe clips were placed into individual 1.5 ml micro-centrifuge tubes containing 1 ml of 70% ethanol. Following sampling, all animals were released back into the breeding ponds. The number of individuals sampled per each population varied between 4 (Witham) and 61 (Mitcham and Palmer’s Green) with a mean of 30 animals sampled per site (Supplemental table S1.).

### Age determination

Age was determined by skeletochronology, following the protocol for *R. temporaria* published by Miaud et al [31] with the following modifications. The phalangeal bone was separated from soft tissues, decalcified with 5% nitric acid for 1.5 hours and washed with water over night. Cross sections (12 µm thick) were then cut from the bone using a cryostat and stained using haematoxylin for 20 minutes. Lines of arrested growth (LAG) were counted using a light microscope at 200x-400x magnification, 10-12 sections were analysed for each individual and two different researchers verified counts. Age at sexual maturity was determined as the youngest age at which inter-LAG space reduced in size, as juvenile inter-LAG space is significantly wider than post sexual maturity [32].

### Statistical Analyses

#### Body size by age and age at sexual maturity

We conducted all statistical modelling in R [33]. We used linear mixed effects regression (lmer) models, implemented in the package lme4 [34] fitted with a Gaussian error structure, and a stepwise simplification procedure to investigate the impact of population ranaviral disease history on the body size (SVL) of *R. temporaria*. Sex, ranaviral disease history of the source population and their interactions were fitted as fixed effects and source population alone as a random effect (Table 1). Since male and female frogs grow at different rates [31,35,36] the datasets of each sex were analysed separately.

**Table 1.** Model summaries of model simplification procedure to evaluate the effect of ranaviral disease history on the body size and age at maturity of *R. temporaria* populations. All models contained only population of origin as a random effect. The p values presented here represent the significance of the parameter removed from the preceding model as calculated by a likelihood ratio test between models (anova in R.) svl = snout to vent length, agemat = age at sexual maturity and df = degrees of freedom of the model.

| Male Body Size |  |  |  |  |
| --- | --- | --- | --- | --- |
| Fixed effects structure | Removed fixed effect | df | Chi<sup>2</sup> | p |
| svl~age * status |  | 6 |  |  |
| svl~age + status | Age * Status | 5 | 0.0587 | 0.8085 |
| svl~age | Status | 4 | 0.21 | 0.6467 |
| svl~1 | Age | 3 | 186.38 | <0.001 |
| Female Body Size |  |  |  |  |
| Fixed effects structure | Removed fixed effect | df | Chi<sup>2</sup> | p |
| svl~age * status |  | 6 |  |  |
| svl~age + status | Age * Status | 5 | 0.9383 | 0.3327 |
| svl~age | Status | 4 | 2.4276 | 0.1192 |
| svl~1 | Age | 3 | 47.16 | <0.001 |
| Male age at maturity |  |  |  |  |
| Fixed effects structure | Removed fixed effect | df | Chi<sup>2</sup> | p |
| agemat ~ status |  | 4 |  |  |
| agemat ~ 1 | Status | 3 | 0.9869 | 0.3205 |
| Female age at maturity |  |  |  |  |
| Fixed effects structure | Removed fixed effect | df | Chi<sup>2</sup> | p |
| agemat ~ status |  | 4 |  |  |
| agemat ~ 1 | Status | 3 | 0.2969 | 0.5858 |

A separate lmer model was fitted to investigate the impact of the ranavirus history status of the source population on age at sexual maturity. In the full model, age at maturity was fitted as the response variable, ranaviral disease history of source population as a fixed effect and source population as a random effect. As female *R. temporaria* mature later than males [31,36] the datasets of each sex were analysed separately.

#### Influence of *Ranavirus* on population age structure

The impact of disease status on the age structure of *R. temporaria* populations was investigated using a Bayesian ordinal mixed effects model implemented in the package MCMCglmm [37]. We fitted age class as an ordinal response variable (9 discrete classes, ages 2 – 10 years), disease status of the source population as the fixed effect and source population as a random effect. We used uninformative priors for both the random effect (G) and residual variance (R) structures, but fixed the residual variance at 1 as this quantity cannot be estimated in ordinal models. The model was run for a total of 600000 iterations with a burn-in period of 100000 iterations and a thinning rate of 500, giving a final sample of 1000 draws from the posterior distributions. Mean probability of membership and 95% credible intervals for each age class for each of the two disease history groups was calculated from the linear predictor. We assessed model convergence using the Gelman-Rubin statistic calculated from two independent chains initiated with overdispersed starting values. All G-R values were <1.05, indication convergence (see supplementary figure S9 for diagnostic plots). Age structure plots (Fig 3) suggest that observed changes were similar for both sexes, so to increase power the dataset was not split by sex.

### Population Matrix Modelling

#### Initial matrix construction and population projections

To investigate the potential for changes in demography to impact the long-term viability of *R. temporaria* populations, we conducted population matrix modelling. An initial female-based Leftkovich matrix was produced consisting of 11 stages; eggs, juvenile (sexually immature frogs) and adult age classes ranging from 2 to 10 years old. Leftkovich matrices are stage based and split the female (providing females are the limiting sex) component of a population into classes based on age, size, developmental stage or similar depending on the study species. The survival of each age class into the subsequent age class is represented as a proportion and the fecundity of each class into every other class (if applicable) as a raw value of females generated per female. See supplementary material for structural examples of Leftkovich matrices. Our basal matrix, representing a putatively disease-free population, was constructed using published vital rates for *R. temporaria* from Biek *et al.* [38]. Biek *et al* assume a constant and uniform maximum reproductive output for all sexually mature adult females of 650 eggs per year. However, it has been shown that the reproductive output of *R. temporaria* is thought to be significantly influenced by body size [38 but 39]. In conjunction with the body size by age data generated in our field study, we adjusted the vital rates for fecundity to increase stepwise for every year post sexual maturity, starting at a rate of 250 eggs for two-year-old breeding animals and increasing by 50 eggs per year until an output of 650 eggs per year was attained at adult age class 10. The rates of Biek *et al.* [38] also assume transition from juveniles directly into adult class 2. However, since fecundity in our matrix is not uniform and female frogs from our dataset were found to mature at ages 2, 3 and 4 (Supplementary Figure S8, consistent with previous findings in *R. temporaria* [31]), we accounted for variation in onset of sexual maturity by allowing juveniles to remain juveniles and to transition directly into adult classes 2, 3 or 4 with highest chance of reaching maturity at adult stage 3. See supplementary table S2 for a numerical representation of the full basal matrix.

We created two age distribution vectors (one for each disease history group) by calculating the proportion of individuals observed in populations of each disease history type that represented each age class and using those proportions as probabilities by which to weight a draw of 150 individual females that could belong to any age class. We used the projection function of the Popdemo package [40] to project these starting populations 20 years into the future based on our starting matrices (additional parameters; standard. A = True, standard. vec = True).

Deaths due to ranavirosis are known to occur annually in diseased populations of UK common frogs [17,24] and annual breeding in permanent water bodies, within which ranaviruses can persist [41], has been identified as a key ecological risk factor of disease [25]. To represent an increased annual chance of an individual dying from ranavirosis every year that it returns to an infected water body, we created additional matrices, which reduced adult survival annually. No age-specific data are currently available on ranavirus prevalence and mortality rates of wild adult amphibians caused by ranavirosis. We therefore modelled the population dynamics of a range of percentage decreases in annual survival in positive disease history populations (1% -25%). We found no significant difference in population dynamics of any starting population based on decreasing annual adult survival alone (supplementary figure S3), we therefore selected a reasonable value of a 5% increased likelihood of mortality per year to represent theoretical populations affected by lethal ranavirosis (Supplementary Figure S3).

#### Stochastic population projections

To investigate the potential impact of environmental stochasticity on populations experiencing persistent disease (5 % increased adult mortality per year of age) and those that are not affected by ranavirosis we created an additional set of matrices to represent differing ecological conditions. A well-known threat to the stability of amphibian populations is climate induced reproductive failure [11,42], and in the UK fluctuations in late winter/early spring night time temperatures can result in frost killing amphibian spawn. Low water temperatures have been identified as the principle threat to *R. temporaria* spawn in UK ponds [43] and are also associated with oomycete infections of amphibian egg masses, which can result in total reproductive failure [44]. To examine the impact of such scenarios, we created alternate versions of both starting population matrices in which fecundity of all sexually mature adult classes was reduced to 0 (supplementary tables S4 and S5).

To examine the impact of potential recurrence of mass mortality events due to ranavirosis, we created a further two matrices; one where the survival of all sexually mature adult classes was reduced to 20%, a biologically plausible but not extreme mortality estimate given that greater than 90% mortality has been observed in mass mortality events caused by ranavirosis [19] and a final matrix representing a catastrophic year in which both adult mass mortality and total reproductive failure occurred (post-maturity adult survival of 20% and fecundity of 0, supplementary tables S6 and S7.) We used the stochastic projection function of the pop.bio package [45] to run the following stochastic projections;

A. Disease-free population with a 10% annual chance of reproductive failure;
B. Population affected by lethal ranavirosis with a 10% annual chance of reproductive failure;
C. Population affected by lethal ranavirosis with a 10% annual chance of either reproductive failure or an adult mass mortality event;
D. Population affected by lethal ranavirosis with a 10% annual chance of either reproductive failure or an adult mass mortality event and a 5% annual chance of both challenges occurring simultaneously.

To ensure all stochastic projection populations started equally, we used the starting population vector of the disease-free populations for all stochastic projections. All projections were run for 100 years and for 5000 iterations. Based on initial population projections we enforced a ceiling carrying capacity for our theoretical populations of 200 sexually mature adult females summed across all age class.

## Results

### Body size by age and age at sexual maturity

We sampled 208 male and 66 female frogs, of which 103 males and 31 females were sampled at populations where lethal ranavirosis had been reported.

For both male and female sub groups, age had a highly significant effect on SVL (males; df = 4, Chisq = 186.38, p<0.001, females; df = 4, Chisq = 47.16, p<0.001; Fig. 2; Table 1). Frogs from positive disease history sites were larger per age class, but this was a non-significant effect (Table 1). Mean age at sexual maturity of males from diseased field sites (n=57) was 2.6 years (± SE 0.07) and 2.8 years (± SE 0.06) at locations where disease was not reported (n=59). Mean age at sexual maturity of females from positive field sites (n=19) was 3.2 years (± SE 0.13) and for females from negative populations (n=17) it was 3.3 years (± SE 0.10), and all differences were non-significant (Supplementary Figure S8; Table 1.)

**Fig 2.**
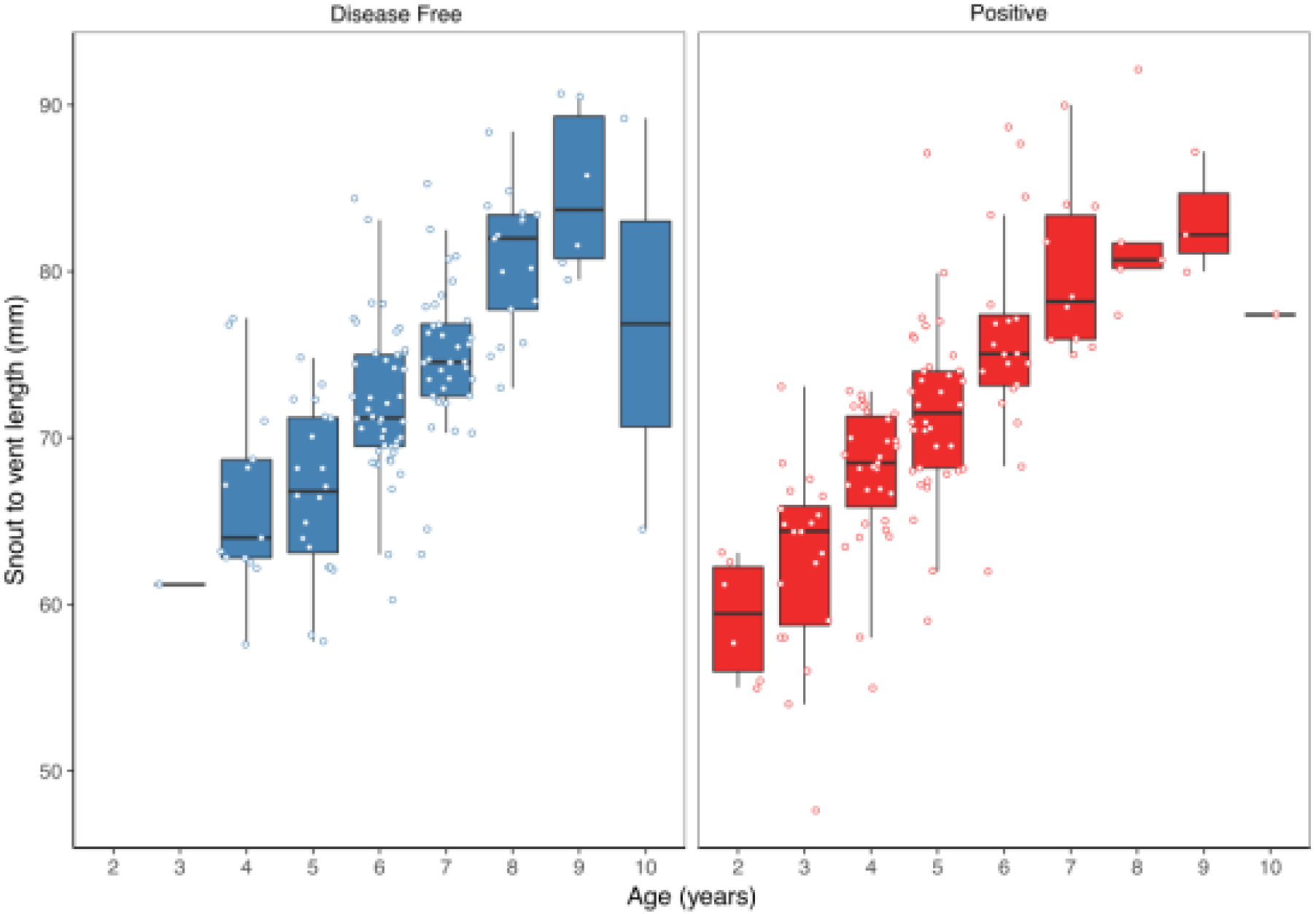
The relationship between snout to vent length (SVL) and age in *R. temporaria* from ranavirus positive and disease free populations. We found no significant effect of disease history on the SVL of frogs. Our data does however show clear evidence of continued growth throughout life regardless of disease history.

### Influence of *Ranavirus* on population age structure

The mean age of males from populations where disease related mortality had been reported was 4.8 years old (± SE 0.16) compared to an average age of 6.3 years (± SE 0.13) at sites where disease has not been reported. Mean female age at diseased sites was 5.3 years (± SE 0.28) compared to 6.6 years (± SE 0.26) at locations where only healthy frogs had been observed.

Disease history had a significant effect on age structure (effect size -1.43, lower 95% credible interval -2.37, upper 95% credible interval -0.38, *p*=0.008; Fig 3), showing populations with a positive history of ranavirosis to be dominated by younger *R. temporaria*. We calculated the difference in posterior probabilities of belonging to an age class based upon disease history status by subtracting the posterior probability of a frog of age X being encountered at positive history population from the posterior probability of a frog of age X being found at a disease-free population. The resulting difference values show that adults aged 2 - 5 years old are more likely to be encountered in disease positive populations, and those aged 6 – 10 years old are more likely to be observed at populations where no disease has been recorded (Fig 4.) Differences in age distributions were strongly supported for all age classes (95% credible intervals (CI) of difference do not cross zero), except for six year olds. Although six-year-old frogs were more likely to be observed in populations where disease had not been reported, the 95% credible intervals incorporated 0 (mean difference = -14.41, 95% CI -30.45 to 3.96; Fig 4).

**Fig 3.**
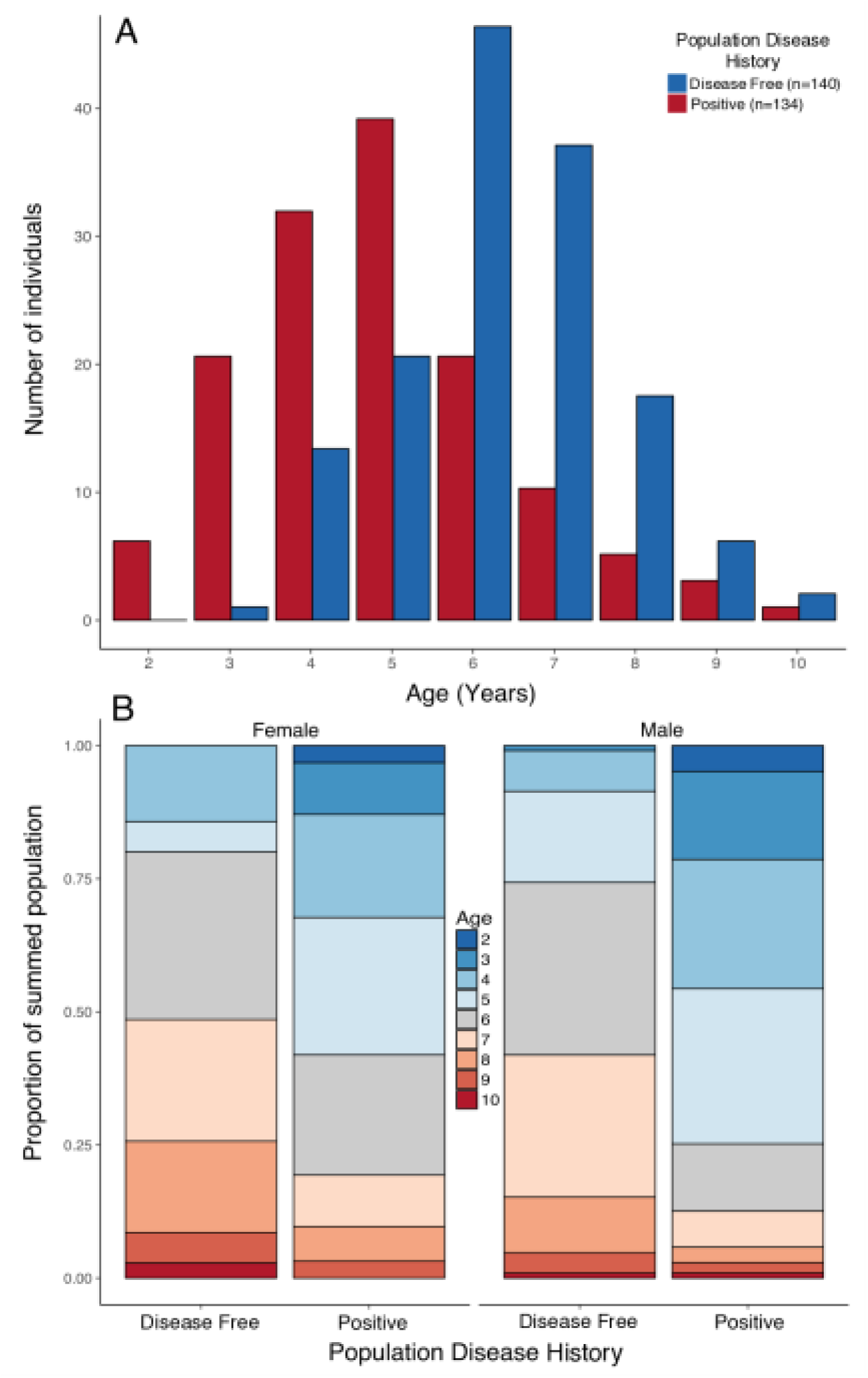
**A)** Histogram of raw counts of numbers of individuals observed per age class per disease history status type. **B)** Proportional stacked bar chart of the proportion of individuals found in populations of each disease history that was a given age, broken down by sex. Breeding populations with a poisitive disease history of ranavirus are dominated by animals 5 years of age and younger. Populations with no history disease are majorly comprised of animals 6 years of age and older.

**Fig 4.**
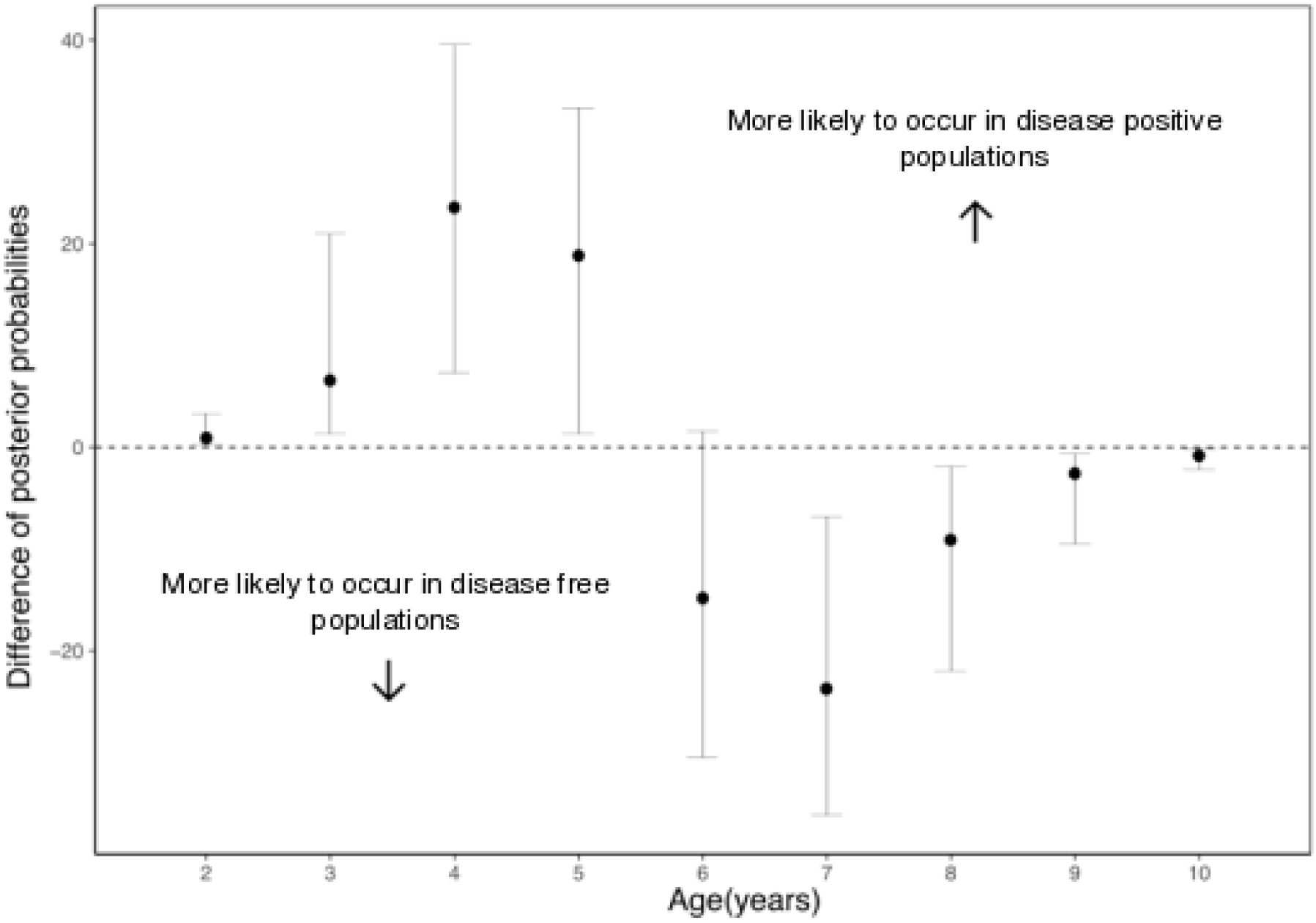
The mean difference in the posterior probabilities of belonging to a given age class by population infection history status. Values > 0 indicate that an age class is more likely to be observed in a positive disease history population and < 0 a putatively disease free population. An age with 95% (2.5% - 97.5%) credible intervals that do not span zero suggests that influence of disease status on that age class is supported by our model. This is the case for all classes other than age 6 which although found to be observed more often in disease free populations has credible intervals spanning 0.

### Population projection modelling

Initial projections of the population dynamics of positive (5% reduction in adult survival per year of age) and disease free populations showed that both followed approximately equal trajectories. Population growth initially spiked due to an influx of pre-mature age classes into the population, followed by a period of attenuating oscillation before reaching equilibrium at around the fifteenth year (Supplementary Figure S10). Final sizes of disease positive populations were non-significantly smaller (positive n= ~146 vs disease free n= ~ 182; t-test, t = 1.67, df = 39.76, p = 0.102).

Disease free populations subject to a 10% chance of recruitment failure per year were still able to consistently attain populations sizes near carrying capacity much more often than any disease positive population (Fig 5, table 2). Increasing complexity of stochastic scenarios reduced the viability of populations experiencing ranaviral disease. Disease positive populations where both adult mass mortality and reproductive failure could occur within the same year were the least viable, being driven locally extinct in 58% of model iterations (Fig 5, table 2).

**Table 2.** The number of iterations per 5000 that each stochastic projection model reached a given population size. Disease Free = Simulated disease free population under a 10% annual chance of complete reproductive failure. Positive = Simulated positive disease history population under a 10% annual chance of complete reproductive failure. Positive A = Simulated positive disease history population under a 10% annual chance of reproductive failure and a 10% annual chance of a recurrent adult mass mortality event in exclusive years. Positive B = Simulated positive disease history population under identical conditions to Positive A with addition of a 5% annual chance of complete recruitment failure and adult mass mortality in the same year. *K* = Imposed population carrying capacity of 200.

| Model | Extinct | <50 | 50-100 | 100-150 | 150-199 | *K* |
| --- | --- | --- | --- | --- | --- | --- |
| **Disease Free** | 12 | 525 | 177 | 734 | 1319 | 2233 |
| **Positive** | 429 | 4062 | 443 | 48 | 15 | 3 |
| **Positive A** | 611 | 4300 | 87 | 2 | 0 | 0 |
| **Positive B** | 2918 | 2081 | 1 | 0 | 0 | 0 |

**Fig 5.**
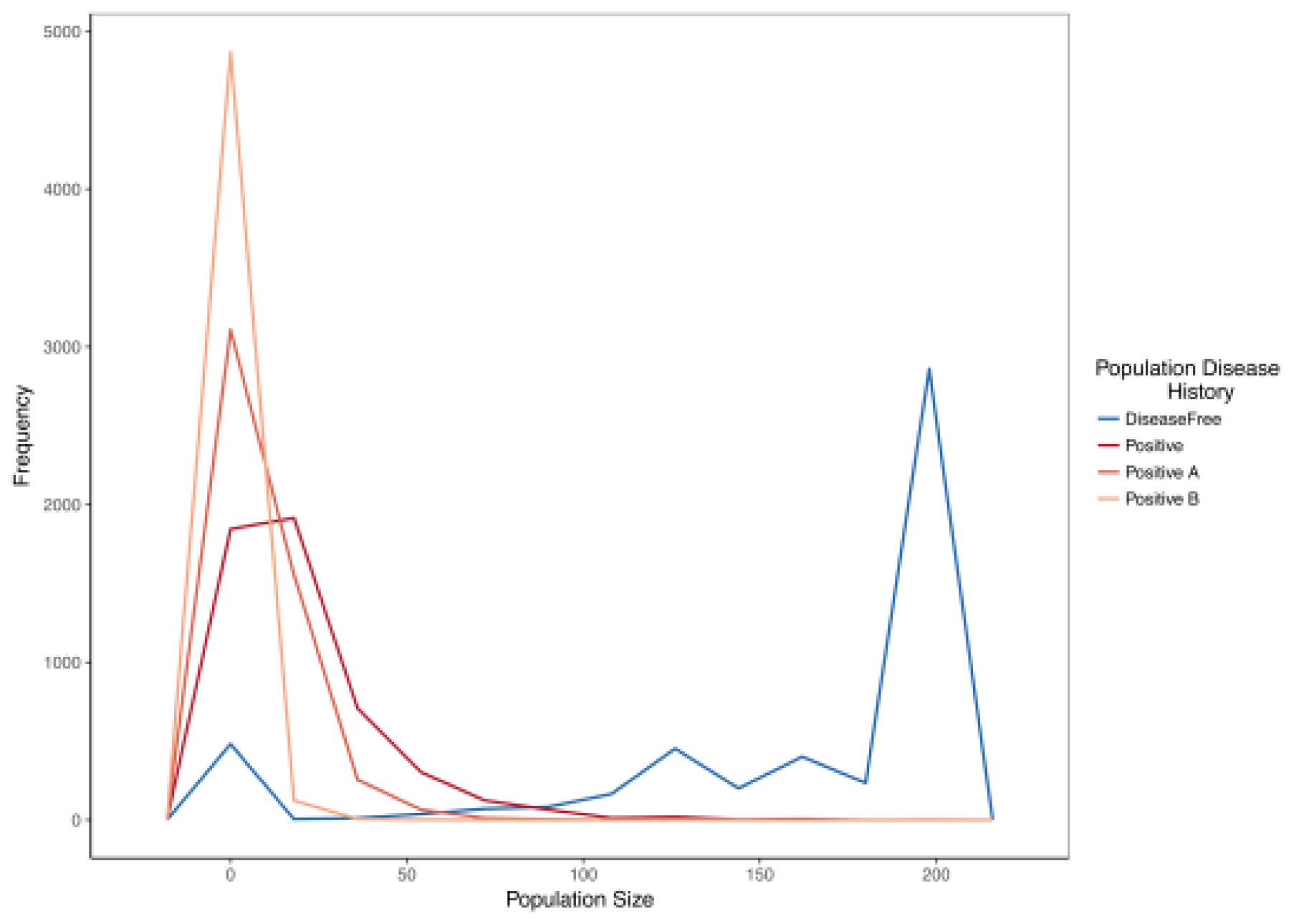
Frequency polygon of iterations in which the projected population hit a given size in stochastic projection modelling. The same starting population vector based on summed observed disease free populations was used in all models. Disease Free = Simulated disease free population under a 10% annual chance of complete reproductive failure. Positive = Simulated positive disease history population under a 10% annual chance of complete reproductive failure. Positive A = Simulated positive disease history population under a 10% annual chance of reproductive failure AND a 10% annual chance of a recurrent adult mass mortality event in exclusive years. Positive B = Simulated positive disease history population under identical conditions to Positive A with addition of a 5% annual chance of complete recruitment failure and adult mass mortality in the same year.

## Discussion

Our results provide the first evidence that a history of ranavirosis in amphibian populations may drive shifts in the age structure of the mature component of host populations, potentially caused by impacts on host life history strategy. We found that breeding populations of *R. temporaria* with a history of ranaviral infection were significantly more likely to contain individuals of 2 – 5 years of age than were breeding populations without a known history of exposure to ranavirus. Conversely, disease free populations were more likely to contain older individuals aged 6-10 years old. Our subsequent population matrix modelling revealed that the demographic differences between disease positive and putatively disease free populations might increase the vulnerability of the former to variable environmental conditions that affect recruitment and adult survival. Our data suggest that challenge by disease can mediate shifts in individual life histories that subsequently increase the susceptibility of populations to declines caused by stochastic environmental conditions. This provides a mechanism explaining why, even in the absence of high rates of mortality in younger age classes, disease-challenged populations can suffer higher rates of extirpation over more protracted time scales.

### The impact of disease on body size and age at sexual maturity

Life history theory predicts that when mortality rates in adults are high, reproductive fitness is maximised when reproduction begins earlier in life [46]. We therefore hypothesised that *R. temporaria* originating from populations where ranavirosis causes increased adult mortality would reach sexual maturity at an earlier age than frogs from disease free populations. We also hypothesised that a trade-off in the allocation of resources to early reproduction rather than growth would cause frogs from disease positive populations to attain a lower body size per age than their counterparts from disease free populations. Both of these hypotheses were also based upon empirical studies conducted on populations of other vertebrates subjected to persistent infectious diseases [1–4]. Surprisingly, we found that neither of these hypotheses were supported by our data. We found that a positive disease history of ranavirosis does not have a significant impact on either; age at sexual maturity or body size throughout life. This is in contrast with the findings of Scheele *et al* [47] who did find that an infectious disease (*Bd*) reduced the size and age at sexual maturity of *L. v. alpina.*

The age at which *R. temporaria* mature is intrinsically linked to attaining a minimum body length need to successfully reproduce [31,35]. The age at which this length is attained has been shown to be heavily influenced by several environmental factors such as photo-period and altitude [31]. All frogs in our study were found to mature at either 2, 3 or 4 years of age and this is consistent with the findings of other similar studies on *R. temporaria* [31,36,35]. In fact, no previous study has found male or female *R. temporaria* to reach maturity younger than 2 years of age and it has been shown that post-sexual maturity there is little detectable trade-off between growth and fecundity in response to sub-prime environments [48]. This suggests that the life history strategy of *R. temporaria* may already be optimised to generate maximum reproductive fitness in light of other factors of their environment and that scope for further plasticity in traits such as age at sexual maturity and subsequent growth rates upon the introduction of an infectious disease may be minimal.

### The impact of disease on population age structure

Our results illustrate that a history of disease can have a significant effect on the age structure of a population. We found that older *R. temporaria* (aged 6 -10 years) are significantly relatively absent within populations that have a positive history of ranaviral disease. *Rana temporaria* are highly philopatric [49], and our study populations occupy permanent, urban or semi-urban garden ponds [24,27]. Ranaviruses have been shown to be able to persist in permanent water bodies [41], particularly in the presence of suboptimal or secondary host species [25,50]. Mortality due to ranavirosis is annually recurrent [17,24] and unlike many similar host-ranavirus systems globally, infection in the UK seems to primarily affect adult life stages of *R. temporaria* [22,51]. Apparent adaptive immune response of *R. temporaria* to ranaviruses is limited [52]. It is therefore possible that adult mortality in infected populations is maintained in such a way that an individual is more likely to become infected and die from ranavirosis the more often it returns to spawn in infected water bodies. Such a scenario, would explain the decreased likelihood of observing animals greater than 5 years of age in populations with a positive disease history of ranavirosis.

Concurrently, we found that frogs aged 2 – 5 years old were significantly more likely to be present in populations that have never shown signs of ranavirus infection. Given the apparent lack of compensatory change in age at sexual maturity, which may cause such a pattern, this clear overabundance of young animals in the breeding ponds at disease free populations is more difficult to explain.

Theory suggests that the first compensatory response to high adult mortality should come in the survival rates of lower age classes [5]. We lack any data on the immature age classes present at our study populations and the snapshot nature of our study means we cannot draw inference on whether the survival rates of younger adult age classes are increased in ranavirus positive populations. However, it is plausible that such an increase could result in the overabundance of younger animals.

Alternatively, the fact that we encountered 2 and 3-year-old breeding frogs at ranavirus positive populations but not disease free populations suggests that the relatively high number of young animals might be the result of behavioural changes that enhance the life time reproductive success of individuals in the face of high adult mortality.

Participation in spawning events in exposed aquatic environments is associated with significant mortality risk to an individual, caused either by exposure to predation or the act of mating itself [44]. Additionally, smaller *R. temporaria* present in breeding populations are easily out competed by larger individuals, who are more likely to secure a mate and achieve amplexus [36]. Participating in breeding events when the chance of reproductive success is low is potentially counter-productive to an individual’s reproductive fitness meaning that small, less competitive individuals may defer breeding attempts until they are older and larger. However, in environments of high adult mortality the number of lifetime reproductive events is potentially limited. In such environments lifetime, reproductive success is likely to be higher when an individual takes advantage of all possible chances to produce offspring. This may result in smaller, less competitive individuals attempting to breed earlier than they would do in the absence of disease induced mortality.

The loss of larger, more competitive individuals at disease positive field sites may also release the smaller individuals from intraspecific competitive pressure. Increased opportunity for less competitive individuals to participate successfully in reproductive events may also result in animals that would normally defer breeding attempts until subsequent years or larger body sizes attempting to reproduce earlier. Any of the above described mechanisms, or a combination of the three would lead to an increase in the number of younger animals in disease positive breeding populations relative to populations that have never experienced disease.

### The potential impact of disease on the viability of populations

When modelled in the absence of environmental variability our representations of both ranavirus positive and disease free populations were able to attain growth rate equilibrium and persist indefinitely. However, the likelihood of extinction increased in simulated positive disease history populations as higher levels of environmental stochasticity were incorporated into our model. These results are consistent with previous observations of age structure truncation increasing the vulnerability of populations to environmental stochasticity [3,11,53].

We are afforded a degree of confidence in our population models as the results corroborate previous empirical data collected from the same study system. A long term study of population sizes has shown that following an outbreak of ranavirosis, characterised by a mass mortality event, UK frog populations follow three possible trajectories. These are: 1) complete recovery to post outbreak population levels, 2) persistence at a largely reduced population size, or 3) local extinction [24]. Our simulated *R. temporaria* populations were projected to population sizes that incorporate all of these possible outcomes, dependant on the levels of environmental stochasticity to which a population was subjected.

Body size, age and fecundity are tightly positively correlated in *R. temporaria* [39] as well as in many other species [e.g. 53–57]. The relative absence of the oldest and largest breeding animals from disease positive populations means that per capita fecundity will be reduced and annual recruitment rates lowered. Such changes likely heighten the impacts of any events that result in failed recruitment or further adult mortality.

Our simulations suggest that the fate of an infected population may depend heavily upon the environmental conditions in the years following an outbreak, providing a potential explanation of why some populations appear to persist in the presence of an infectious disease, while others are driven to local extirpation. Although we have demonstrated the importance of environmental variation in on the within population disease dynamics of ranavirus, it is not thought to play a role in the wider context of disease spread [27]

### The importance of variations in fecundity in population models

It is well known that *R. temporaria* grow continually throughout life [31,35,48], and our data show this to be true in our study populations. We therefore took variations in fecundity by body size into account when constructing our population projection matrix models. In the absence of increased extrinsic mortality, populations simulated by a matrix with varied fecundity were stable and viable. However, when increased adult mortality due to ranaviral exposure was incorporated into our models, the lower fecundity of younger adult age classes played a role in the reduced viability observed. Variations in adult fecundity due to body size are often not included in population projection modelling [eg. 60 but see 61]. However, this result highlights the importance of considering age or body size specific changes in fecundity in population modelling, particularly when considered threats disproportionately impact certain age classes. Ensuring matrix models of populations represent the life history of the study species as closely as possible is essential, especially when seeking to inform conservation or policy decision making.

## Conclusion

Empirical evidence is beginning to accumulate that infectious disease can alter the demographic structure of populations. This is of particular concern in species that are vulnerable to extinction. Our results provide the first evidence that infection with a ranavirus results in age truncation of amphibian populations, despite no obvious change in age or body size at sexual maturity. Population projections showed that the impact of age structure truncation on disease positive populations could potentially be substantial, increasing vulnerability to environmental fluctuations that affect recruitment success. Our results highlight an increasing need to better understand the impact of disease on life history and the demography of host populations. Further investigation of this relationship, possibly via a long-term mark-recapture study on the same populations used here could elucidate the exact mechanisms by which these demographic shifts are generated. This work provides evidence that the emergence of an infectious disease within a population can heighten its vulnerability to external stressors. Although in this case our theoretical stressors were environmental in origin the same is likely to be true for all types of stressor, including anthropogenic.

## Ethics statement

This project was approved by the ethics boards of both the University of Exeter and Zoological Society of London and conducted under the project license 80/2466 issued by the UK Home Office. All field sampling was conducted under the personal Home Office license 30/10730 issued to Lewis Campbell.

## Data accessibility

The raw data file and R scripts used in the analyses of this work are available online via GitHub at the following link;

https://github.com/zoolew/Ranavirus-FrogDemography

## Author contributions

LJC devised the study, collected field samples, prepared samples in the laboratory, analysed data and wrote the manuscript. GT performed skeletochronology and provided editorial comment on the manuscript. TWJG devised the study, provided important logistical support and editorial comment on the manuscript. BCS provided vital input in the analyses of the data and editorial comment on the manuscript. AGFG and LW provided significant advice throughout and editorial comment on the manuscript. XAH devised the study, analysed the data, and provided significant editorial comment on the manuscript.

